# The Quick and the Dead: Microbial Demography at the Yeast Thermal Limit

**DOI:** 10.1101/055442

**Authors:** Colin S. Maxwell, Paul M. Magwene

## Abstract

The niche of microorganisms is determined by where their populations can expand. Populations can fail to grow because of high death or low birth rates, but these are challenging to measure in microorganisms. We developed a novel technique that enables single cell measurement of age-structured birth and death rates in the budding yeast, Saccharomyces cerevisiae, and used this method to study responses to heat stress in a genetically diverse panel of strains. We find that individual cells show significant heterogeneity in their rates of birth and death during heat stress. Genotype-by-environment effects on processes that regulate asymmetric cell division contribute to this heterogeneity. These lead to either premature senescence or early life mortality during heat stress, and we find that a mitochondrial inheritance defect explains the early life mortality phenotype of one of the strains we studied. This study demonstrates how the interplay of physiology, genetic variation, and environmental variables influences where microbial populations survive and flourish.

## Introduction

Population growth rate is arguably the most measured variable in all of microbiology. It is used to study the effects of genetic mutations (Breslow et al. 2008), as an indicator of physiological stress (Warringer et al. 2003), as a readVout of selection (Vasi, Travisano, and Lenski 1994), and as a proxy for fitness (Joseph and Hall 2004). However, net population growth rate is itself a composite function of the birth and death rates of individuals and measuring only the population growth rate confounds them. Furthermore, birth and death rates frequently change with the age. Because analyses of population growth play such a critical role in studies of microbes, disentangling the effects of birth, death, and aging on population growth should lead to a better understanding of microbial physiology, ecology, and evolution.

The range of environmental conditions where the population growth rate of a species is positive corresponds to what the ecologist G. E. Hutchinson called a species' fundamental niche (Hutchinson 1957). The limits of a species' fundamental niche are determined by a combination of physiological, genetic, and demographic factors. One strategy for identifying specific factors that limit the environments where an organism can live is to dissect the demographic mechanisms that drive decreases in population growth rate when a population is exposed to conditions near those limits. Is a decrease in population growth rate driven by a population wide decline in reproductive output as environmental conditions approach their limit? Alternately, are there increases in mortality, perhaps in an ageVstructured manner, that limit growth? How do such environmentally induced shifts in demographic parameters vary with genetic background? Questions like these can only be addressed by characterizing the fecundity, mortality, and age of individuals in populations. Such individual level characterization is particularly challenging in microbes due to their small size, large populations, and the consequent challenges of tracking or recovering individual cells.

Temperature is an environmental variable that plays a critical role in limiting the fundamental niche of microorganisms. Although the total temperature range over which microbial life is found is quite large (at least v5°C to 113°C, (Pikuta, Hoover, and Tang 2008)) most species are confined to a relatively modest range of temperatures that support robust growth. The ability to maintain a relatively high growth rate at high temperatures is thought to be critical to the ecological success of the budding yeast *Saccharomyces cerevisiae. S. cerevisiae* is able to grow at higher temperatures relative to closely related species (Salvado et al. 2011; Warringer et al. 2011). ThermoVtolerance also varies widely between strains of *S. cerevisiae* (Liti et al. 2009) and has a complex genetic basis (Steinmetz et al. 2002; Sinha et al. 2006). This genetic diversity for growth at high temperatures may also affect human health; a high thermal limit has been hypothesized to contribute to the opportunistic invasion of human hosts by *S. cerevisiae* (McCusker et al. 1994).

Using a newly developed method, called TrackScar, that allows us to simultaneously measure the birth rate, viability, and age of individual cells, we analyzed a genetically diverse panel of yeast strains during growth near their upper thermal limits. We find that disparate demographic and physiological responses limit population growth of different strains during heat stress. Furthermore, the ageVstructure of mortality can differ with genetic background, leading to both “premature senescence” as well as “early life mortality.” We show that in one genetic background early life mortality likely results from a temperature sensitive mutation affecting mitochondrial inheritance. By measuring the demographic rates of individuals, we elucidate the diverse physiological and demographic factors that limit the niche of budding yeast.

## Results

### TrackScar measures the fecundity and mortality of single cells

During the process of asexual mitotic reproduction (budding) in *S. cerevisiae* and related yeasts, ringVshaped scars of chitin are formed on the cell wall at the site of cytokinesis in both mother and daughter cells. These rings, known as “bud scars”, provide a cellular record of the number of daughters an individual cell has produced. Bud scars can be visualized using fluorescently labeled compounds that preferentially bind to chitin (Pringle 1991). Wheat germ agglutinin (WGA), a lectin, is a particularly suitable bud scar stain because it can be can be conjugated to a variety of fluorophores and has low background staining of the cell wall.

We developed an experimental method called “TrackScar” for measuring the reproductive output (fecundity) of individual yeast cells based on the sequential staining of bud scars (Fig. 1a,b). The principle behind TrackScar is simple: 1) label a population of yeast cells at two time points (t_0_, *t*_t_), using different fluorophores at each time (e.g. cyan at *t*_0_, yellow at t-J; 2) image cells that were stained at both time points; and 3) quantify the number of cell divisions individual cells have undergone in the given time interval by counting the difference in the number of bud scars produced (e.g. no. of yellow scars-no. of cyan scars). The number of daughter cells produced in a given time interval is a discrete and unambiguous measure of the rate of fecundity at the single cell level. Detailed experimental procedures are provided in the supplementary information.

**Figure 1.**
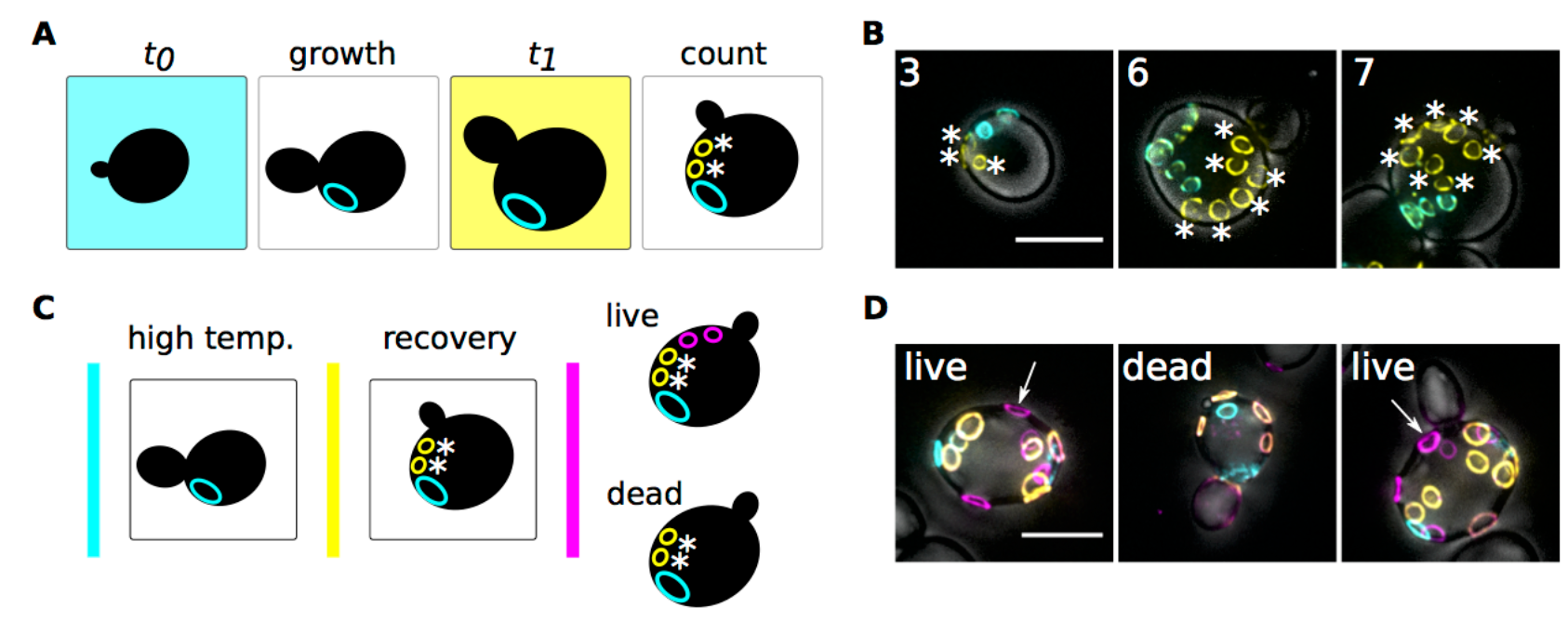
TrackScar measures the fecundity and viability of individual yeast. **A**) A schematic of twoVcolor TrackScar is shown. To measure the reproductive output of individuals, cells are: 1) stained with wheat germ agglutinin (WGA) conjugated to Alexa488 (cyan) at *t*_0_; 2) grown in the absence of the stain; 3) stained with WGA conjugated to tetramethylrhodamine (yellow) at *t_1_.* New buds (labeled with asterixes) are only stained by the second dye. **B**) Examples of haploid S288C cells that produced different numbers of daughters during 6 hr of growth at 30°C in YPD are shown. **C**) A schematic of threevcolor TrackScar is shown. Two colors of WGA (cyan and yellow) are used to measure the fecundity of cells during heat stress. To determine viability, the cells are transferred to permissive conditions and then stained with a third color of WGA (magenta). Viable cells produce new buds that are only stained by the third dye. **D**) Examples of diploid YJM693 cells that were either viable or inviable after exposure to growth at 35.5°C are shown. Arrows highlight one of the bud scars produced by live cells in permissive conditions. All micrographs are maximum intensity projections of zvstacks. Scale bar shows five microns.

Exposing cells to stressful conditions is often accompanied by a decrease in fecundity, with extreme stress leading to cell death. Since twoVcolor TrackScar only measures the number of daughters that a cell produces in one interval, it is possible that cells with low observed fecundity could have died before the first stain (WGA stains both live and dead cells), or in the interval between stains. In order to distinguish between slowly dividing but still viable cells and cells that are dead, the TrackScar method can be extended to include a “recovery phase” (in this case six hours) under permissive growth conditions (30°C), followed by staining with a third fluorophore. Assuming the recovery phase is sufficiently long, cells that show at least one reproductive event during the recovery phase are still viable; cells that show no growth during recovery are either dead or severely growth arrested (Fig. 1c,d). ThreeVcolor TrackScar provides more information about individual cells, but data collection is more difficult because mother cells are further diluted during the recovery phase and because more bud scars must be counted.

We performed control experiments to demonstrate that: 1) TrackScar is sensitive; 2) that staining cells using WGA does not affect their division rate; 3) and that six hours of recovery is sufficient to distinguish live from dead cells. These results are described in the Supplementary Information.

### Identifying diverse responses to heat stress

The fundamental niche of a species is limited by the range of temperatures (‘thermal limits’) in which it can survive. Within a species the thermal limit can vary substantially, reflecting genetic diversity. To disentangle the demographic basis of heat sensitivity we applied TrackScar to a genetically diverse panel of *S. cerevisiae* strains growing in a range of thermal regimes.

#### Identifying slow growing yeast populations

We estimated the maximum population growth rates for a panel of 93 sequenced yeast strains (Strope et al. 2015) across a range of temperatures from 21°C to 40°C by measuring the timeVvarying optical density of liquid cultures grown in a spectrophotometer (Fig. S3). At low temperatures, we observed only small differences between genetic backgrounds in their population growth rates. However at 35.5°C and above there is substantial strainVtoVstrain variation.

To identify heat sensitive versus robust strains, we calculated the ratio of maximum growth rates at 35.5°C, a potentially stressful temperature with high betweenVstrain variance, relative to 30°C, a standard permissive temperature. Strains with a thermal growth ratio near 1.0 were considered robust to thermal stress, while those with ratios significantly lower than 1.0 were classified as heat sensitive (Fig. 2a).

**Figure 2.**
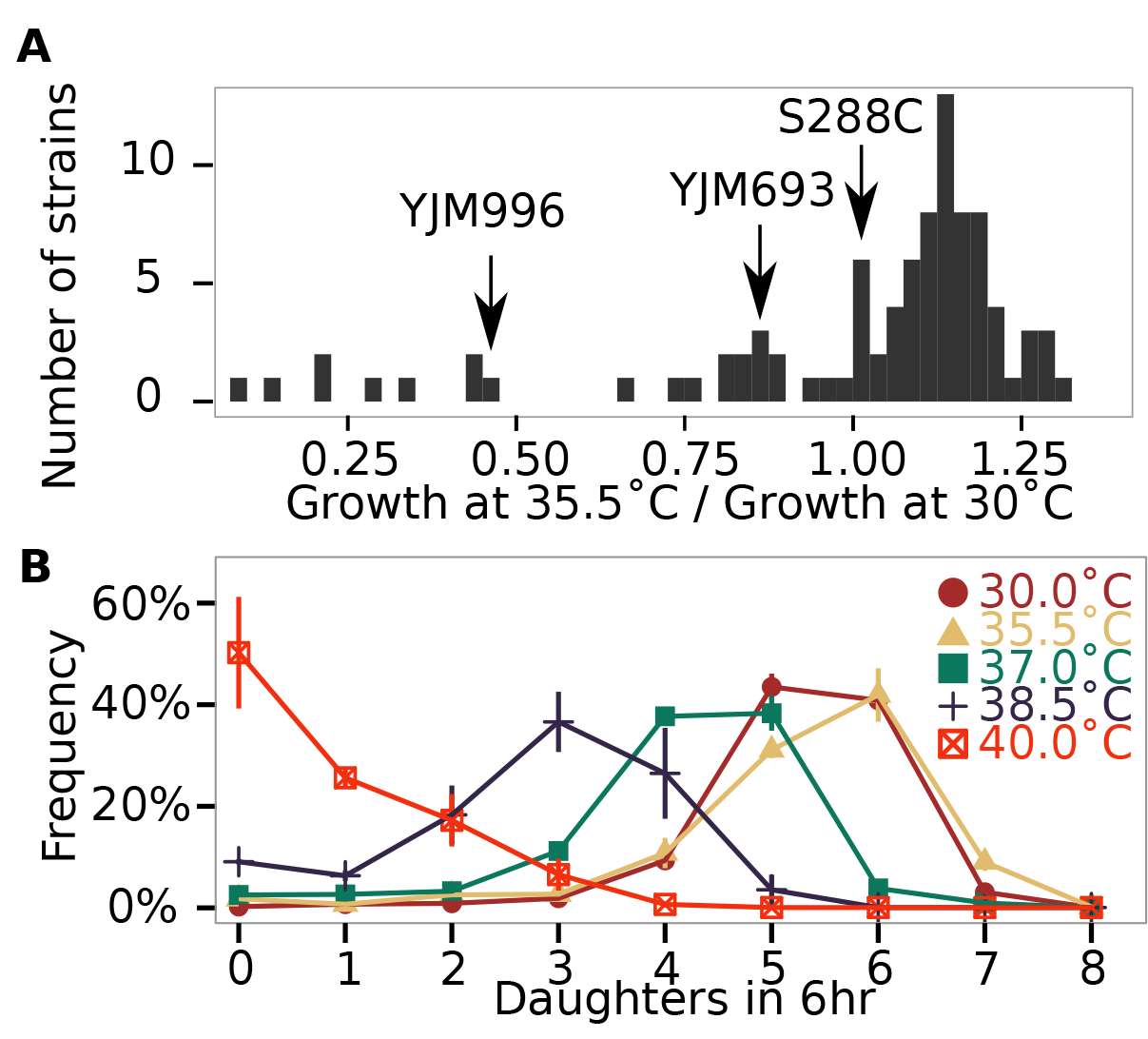
There is extensive genetic diversity in thermotolerance among environmental yeast isolates. **A**) A histogram of the growth rate of each of 93 yeast strains at 35.5°C normalized to their growth rate at 30°C is shown. **B**) Frequency polygons of the number of daughters produced by cells in the strain S288C during 6 hr of growth at 30°C (brown circles), 35.5°C (gold triangles), 37°C (green squares), 38.5°C (black crosses), and 40°C (red crossed boxes) are shown. Error bars are the standard error of the mean for three biological replicates.

TrackScar provides a richer view of how population fecundity changes across environments than measuring only the population growth rate. For example, Figure 2b illustrates how the population distributions of fecundity change over a temperature range of 30°C to 40°C for diploid S288c cells. S288c grows well at both 30°C and 35.5°C. By contrast, at higher temperatures the distribution of fecundities shifts left and there is a noticeable increase in the variance.

### Heat sensitivity is associated with the emergence of heterogeneous subpopulations

We examined the distributions of fecundity for strains that were both sensitive and robust to heat stress. At 30°C most strains exhibit unimodal and approximately symmetric fecundity distributions (Fig. S4), although the mean and variances of fecundity differ between backgrounds. At 35.5°C robust strains continue to exhibit symmetrical fecundity distributions, with only modest differences in mean fecundity. By contrast, many sensitive strains (Fig. 3; Fig. S4) have asymmetric fecundity distributions, with a substantial increase in slow growing or nonVgrowing cells, even leading to bimodal distributions (Fig. 3; Fig. S4).

**Figure 3.**
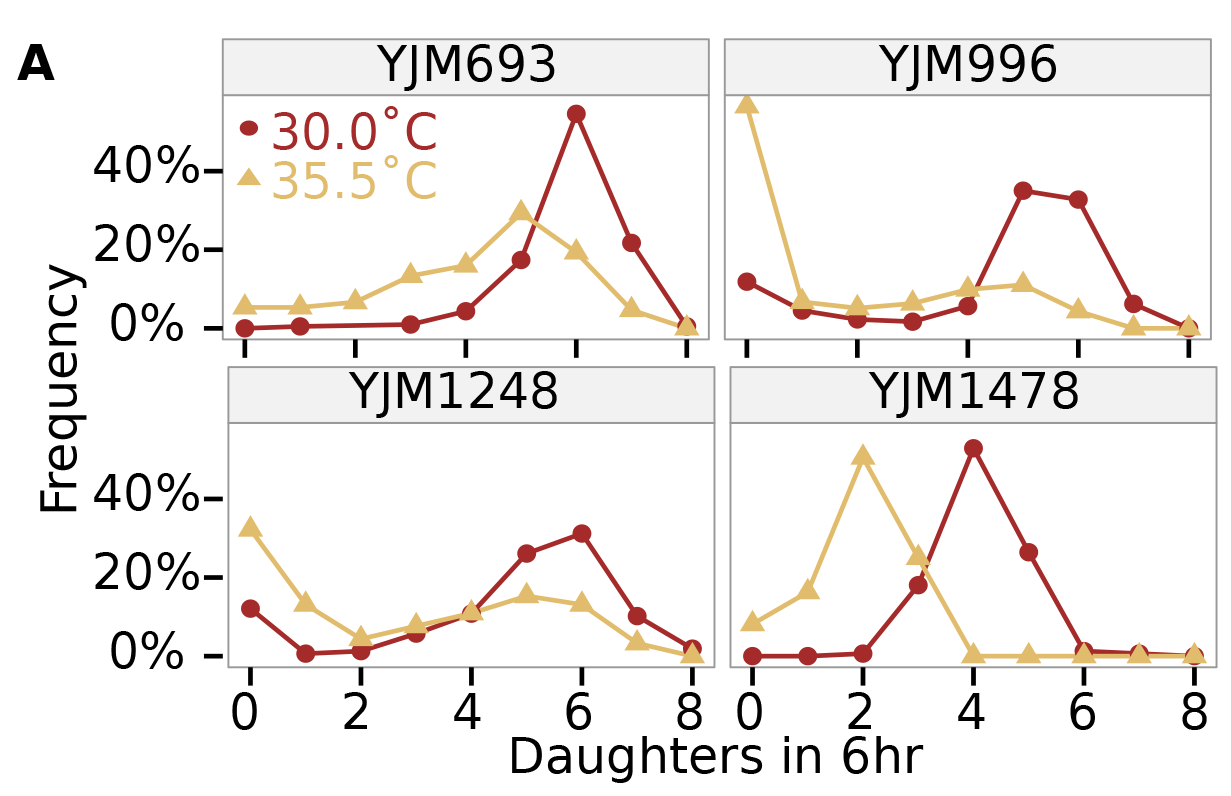
Strains that are sensitive to heat stress frequently have subpopulations with very low fecundity. Plots showing frequency polygons of the number of daughters produced in 6hr at 30°C (brown circles) and 35.5°C (gold triangles) are shown for A) robust and B) sensitive strains are shown. A single representative replicate is shown for each temperature for each strain.

### Increased mortality explains slow population growth for some heat sensitive strains

One mechanism that could lead to asymmetric and bimodal fecundity distributions we observe during heat stress is if some cells died before or during the interval in which we observe their fecundity. To explore this possibility, we used the threeVcolor TrackScar assay to test whether cells with low fecundity during heat stress were viable when the stress is removed. We focused our analyses on three strains-S288C, YJM693, and YJM996-that have distinct fecundity distributions near their thermal maxima (Fig. 2b, Fig. 3). Since S288C does not exhibit a growth rate decrease at 35.5°C, we assayed this strain at 40°C, a temperature where it exhibits a distinct decrease in fecundity and a large number of nonVdividing cells. The other two strains were assayed at 35.5°C. We measured fecundity for six hours during heat stress, incubated the cells overnight at 4°C, then transferred the cells to 30°C and measured the number of divisions during a subsequent sixVhour period.

We used threeVcolor TrackScar to determine whether increased mortality explained the low observed fecundity of cells during heat stress. We compared the number of daughters produced *during heat stress* of cells that either survived or died during heat stress with the fecundity of unstressed cells. S288C cells with low fecundity during heat stress are frequently alive at the end of the period of stress (Fig. 4). In contrast, viable YJM996 cells frequently produce as many daughters as unstressed cells, but with a broad distribution. Remarkably, and in contrast with the other two strains we examined, the distribution of fecundity among YJM693 cells that survived heat stress was identical to that of unstressed cells (KolmogorovV Smirnov test; p = 0.64). This indicates that heat stress causes YJM693 cells to split into a population of quickly dividing viable cells and another population of unviable cells. Taken together, these results show that in some genetic backgrounds mortality can be the sole cause of slow population growth during heat stress, while in other strains a combination of increased mortality and reduced individual fecundity occurs.

**Figure 4.**
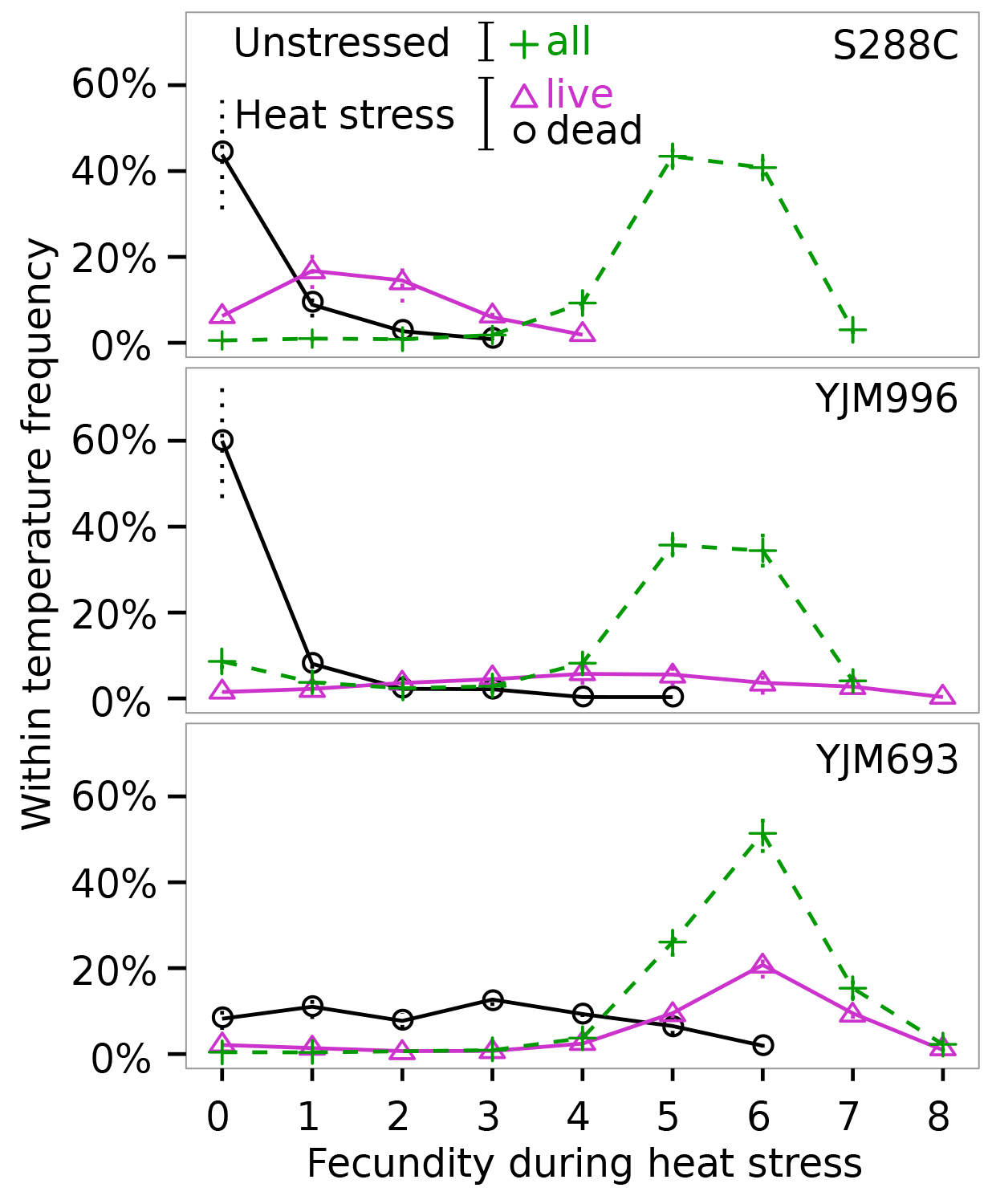
Mortality contributes to slow population growth in a genotype specific manner. Cells were heat stressed at either 35.5°C or 40°C, then assayed for viability at 30°C. The frequency polygons plot the fecundity during heat stress for cells that were subsequently classified as viable (magenta triangles) and inviable (black circles) from strains S288C, YJM996, and YJM693. The fecundity of unstressed cells is plotted as a dashed line with green crosses. Error bars are the standard error of the mean for three biological replicates.

### Fecundity can be positively or negatively associated with age during stress

Yeast mother cells faithfully segregate subVcellular components to their daughter cells, but also asymmetrically retain aging factors such as damaged proteins and mitochondria (Lai et al. 2002) (McFalineVFigueroa et al. 2011; Aguilaniu et al. 2003). Genetic variation could lead to temperature sensitive mutations that affect either of these processes, which could lead to lower fitness in either younger or older cells, respectively. Since TrackScar works by measuring the replicative age of a cell at the beginning and end of an experiment, the age structure of fecundity emerges naturally during analysis. We will call a group of cells with the same replicative age at the beginning of the experiment a “cohort.” To gather sufficient data about older cohorts, we sought out and imaged old cells rather than imaging random cells (see methods).

We measured the average fecundity of cohorts of cells in the strains YJM693, YJM996, and S288C. To test whether fecundity was affected by age, we excluded the fecundity of daughter cells because the fecundity of daughters is lower than mothers due to an extended G1 phase of the cell cycle (Hartwell and Unger 1977)(Fig. S1b). At 30°C, the average fecundity of cohorts of YJM693 increased slightly with age, whereas the fecundity of cohorts of YJM996 was not significantly affected by age (Fig. 5a)(linear model, p = 0.016 and p = 0.68, respectively). The average fecundity of a cohort of S288C was not affected by its age at any temperature (Fig. S5). In contrast, at 35.5°C, replicative age significantly affects the average fecundity of a cohort in the strains YJM693 and YJM996 (linear model, *p* = 0.0006 and *p* = 0.0008, respectively). Interestingly, while YJM693 cells produce an average of 0.21 ± 0.12 *fewer* daughters in six hours per cohort when heat stressed, YJM996 cells produce an average of 0.33 ± 0.17 *more* daughters in six hours per cohort (intervals are 95% confidence intervals of the mean)(Fig. 5a).

**Figure 5.**
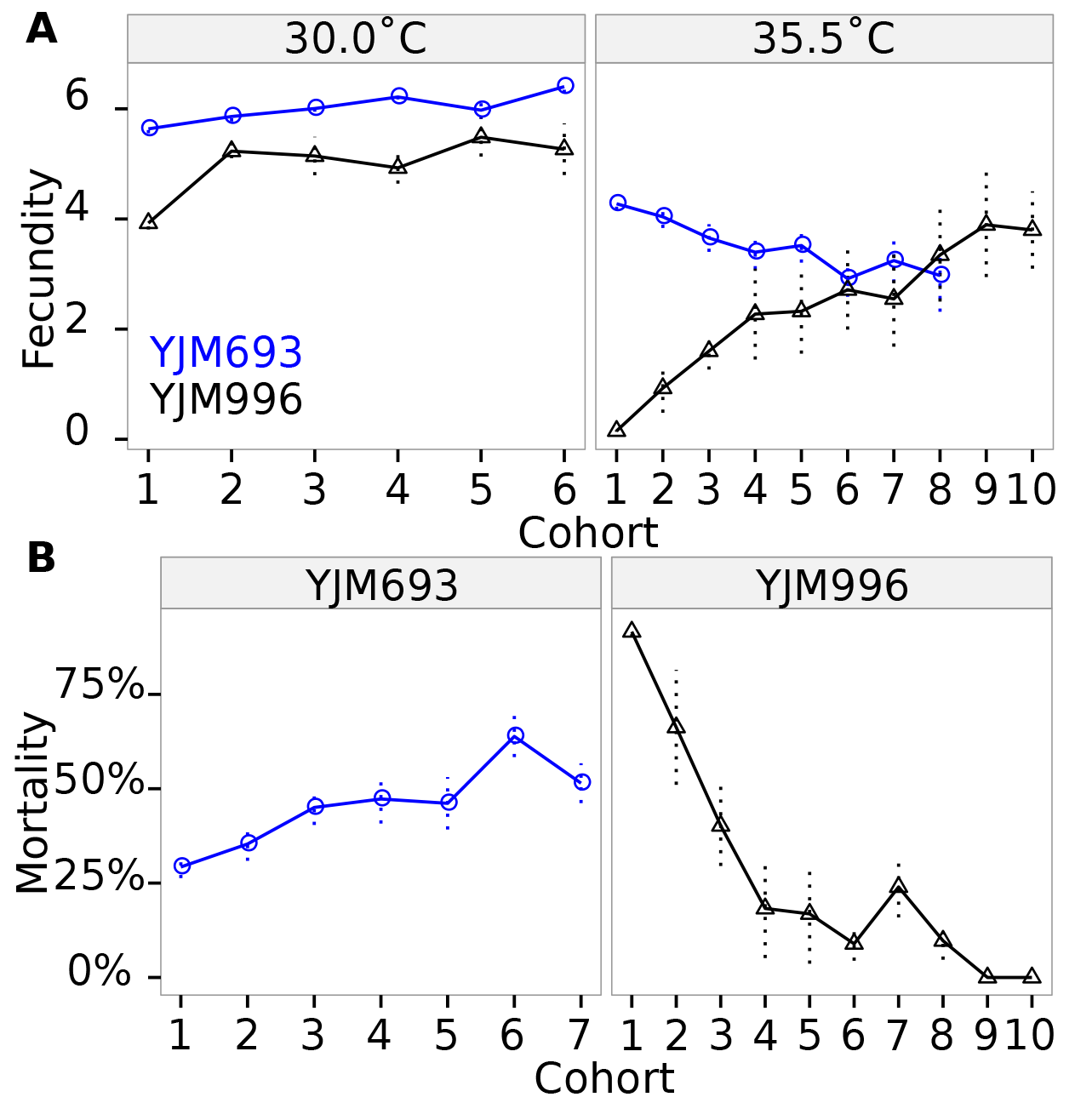
Genetic background determines the age structure of fecundity and mortality during heat stress. A) The average fecundity of a cohort during 6hr grow at either 30°C or 35.5°C for the strains YJM693 (blue circles) and YJM996 (black triangles). Error bars are the standard error of the mean for at least three biologica replicates. Only cohorts with at least three cells in each of the three biological replicates are shown. B) The mortality of cohorts of YJM693 and YJM996 cells is shown. YJM693 cells were considered “dead” if they divided four times or fewer growth at 35.5°C, whereas YJM996 cells were considered “dead” if they did not divide at all during six hours of growth at 35.5°C. Error bars show the standard err of the mean for at least three biological replicates. Only cohorts with at least five cells in each of the three biological replicates are shown.

### Heat stress can cause premature senescence or early life mortality

The fecundity of YJM693 cells is lower in older cells (Fig. 5a). YJM693 cells that divided less than four times during a sixVhour period of heat stress were dead by the end of the experiment (Fig. 4). Consistent with this, we found that the older a cohort of cells was, the more cells in that cohort failed to divide at least four times in six hours-indicating they had died (Fig. 5b). Since, by definition, each cell in a cohort had survived at least long enough to enter the cohort, this indicates that the likelihood of death increases with replicative age at 35.5°C in YJM693 and cannot be explained by a model of constant survival risk. Using logistic regression, we estimate that there is a 21% (95% CI ± 6%) increase in the probability of death for each additional unit of replicative age in this strain during heat stress. These results suggest that thermal stress affects an asymmetrically inherited aging factor in this genetic background.

In contrast to our findings for YJM693, the fecundity of YJM996 cells is higher in older cells (Fig. 5a). Although not all cells with low fecundity in YJM996 died during the exposure to stress, cells that produced no daughters during the six hours of growth at 35.5°C were viable less than 5% of the time (Fig. 4). Therefore, we considered these cells dead for the purpose of estimating ageVstructured mortality. Younger cohorts in the YJM996 genetic background have substantially higher mortality than older cohorts. More than 80% of the youngest cells did not produce a daughter cell during 6 hours of recovery at 30°C, which strongly suggests that they had died (Fig. 5b). Cohorts that had divided only one or two times also had substantially elevated mortality. In contrast, cells in cohorts six and above showed substantially reduced mortality (Fig. 5b). YJM996 thus exhibits an early life mortality phenotype during heat stress. Interestingly, the high mortality of young cells leads older cells to accumulate in the population (Fig. S6). This pattern of mortality in young cells in consistent with heat stress impacting the segregation of a critical cellular component.

### Early life mortality is associated with a failure to inherit mitochondria

The early life mortality phenotype of YJM996 during heat stress suggests that this strain may have a temperatureVsensitive defect in the segregation of some cellular component to daughter cells. Previous studies have shown that failure to properly segregate mitochondria (e.g. in an *Δmmrl* strain) leads to sub-populations of shortV and longVlived cells (McFalineVFigueroa et al. 2011), which could generate mortality patterns similar to the ones in YJM996. We examined the morphology of mitochondria at both 30°C and 35.5°C in strains YJM693, YJM996, and S288C using a mitochondrially localized GFP expressed from a low copy number plasmid. Since this plasmid was uracil selectable, we used *Aura3* derivatives of these strains and grew them in synthetic media lacking uracil.

The strain YJM996 has a temperature sensitive defect in mitochondrial morphology and inheritance. In S288C and YJM693, at both 30°C and 35.5°C, the mitochondria form a branching, threadVlike network. In contrast, the mitochondria of YJM996 grown at 35.5°C formed large globular clumps (Fig. 6). This globular morphology was also present with low (∼10%) penetrance at 30°C (Fig. S7). We noticed that the buds of YJM996 cells often lacked mitochondria even near the end of mitosis (Fig. 6), suggesting that the mitochondria of this strain may be inherited inefficiently. Consistent with this, at 30°C and 35.5°C, approximately 10% and 75%, respectively, of YJM996 cells lack mitochondria entirely (Fig. S7). We observed cells lacking mitochondria to contain nuclei (Fig. 6) and vacuoles, suggesting that the inheritance of other organelles is not affected. Since mitochondria cannot be created *de novo* and must be inherited from mother cells, this indicates that mitochondrial inheritance is inhibited by high temperatures in YJM996.

**Figure 6.**
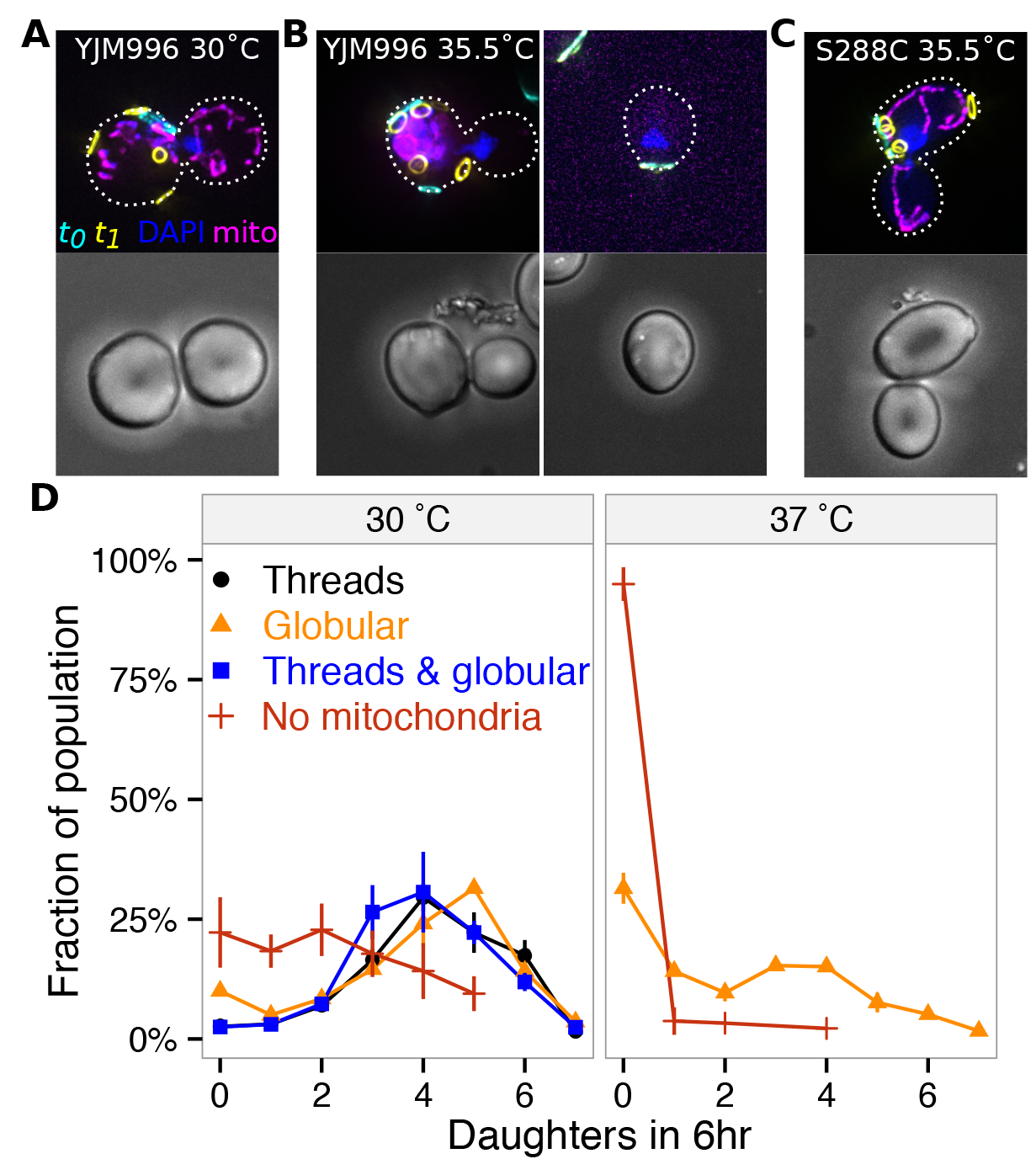
Early life mortality is associated with a defect in mitochondrial inheritance in the YJM996 genetic background. Representative images are shown of TrackScar staining at zero hours (cyan) and six hours (yellow), DAPI staining (blue), and mitochondria-localized GFP (magenta) for a *Aura3* derivative of the strain YJM996 at A) 30°C and B) 35.5°C, and a *Aura3* S288C derivative at C) 35.5°C. Each budding cell is staged in the process of nuclear division to facilitate comparison of mitochondrial inheritance. Dotted lines show the outlines of the cells. D) The fecundity of cells with thread-like mitochondrial morphology, globular morphology, a mixture of clumps and threads, and cells that lack mitochondria is shown for cells in the YJM996 genetic background at both 30°C and 35.5°C. Error bars show the standard error of the mean for three biological replicates of at least 150 cells.

We hypothesized that this mitochondrial inheritance defect was the cause of the early life mortality phenotype in YJM996. Consistent with this, we found that YJM996 cells that lack mitochondria divide fewer times than cells that contain mitochondria at both 30°C and 35.5°C (Fig. S8). Globular mitochondria were found with low frequency at 30°C, and we hypothesized that these mitochondria would be associated with slower growth. However, cells with globular mitochondria produced the same number of daughters as cells with wild-type mitochondria, whereas cells without mitochondria produced few or no daughters at either temperature (Fig. S8). This indicates that mortality in this strain is not caused by a defect in mitochondrial morphology *perse*, although the altered morphology of the mitochondria could contribute to the defect in inheritance. Taken together, these results strongly suggest that YJM693 grows slowly at elevated temperatures due to a defect in the segregation of mitochondria, leading to high mortality in young cells.

## Discussion

We have dissected the demographic mechanisms that delimit the fundamental niche of genetically diverse *S. cerevisiae* strains. The growth rate of a population of microbes in the laboratory at a particular temperature can accurately predict the actual environments (the niche) in which a microbe can thrive (Madigan et al. 2015). However, the growth rate is the sum of the rates of birth and death, and the effect of temperature on growth rate could be due to a change in either. We developed TrackScar to measure age-structured rates of birth and death in populations of budding yeast without the need for trapping or microscopic tracking of cells. TrackScar simplifies experimental design and opens the door to studies of microbial demography in environments inaccessible to microscopic tracking of individuals.

SingleVcelled organisms rarely behave identically, even if they share the same genome and the same environment. Random events like stochastic gene expression can cause such differences (Spudich and Koshland 1976; Maamar, Raj, and Dubnau 2007; Bumgarner et al. 2012). However, predictable events such as cell cycle and aging (Avery 2006; Levy, Ziv, and Siegal 2012), and epigenetic inheritance of growth rate (Ziv, Siegal, and Gresham 2013; Levy, Ziv, and Siegal 2012) can also contribute. Genetic variation can impact cellular individuality, indicating that it could be selected for during evolution (Ziv, Siegal, and Gresham 2013; Fehrmann, BottinV Duplus, et al. 2013; Holland et al. 2013). This individuality makes the average a poor predictor of the behavior of individuals, and can obscure the physiological basis of populationVlevel phenotypes.

Heat stress often causes extensive variability in the fitness of individual cells, as shown by the asymmetric and frequently bimodal distributions of the fecundity we observed. This variability can be age structured, and in at least one case is a consequence of the failure of cells to appropriately segregate subVcellular components (see below). The ability to easily distinguish mother and daughter cells in budding yeast make these patterns particularly apparent, but subtle asymmetries also lead to aging in ostensibly symmetrically dividing organisms such as *Escherichia coli* (Stewart et al. 2005). We speculate that genetic variation in cellular mechanisms that establish or normalize asymmetries between mother and daughter cells may be an important cause of phenotypic variability in other organisms.

Active processes ensure that sufficient quantities of organelles and other subVcellular components are segregated to daughter cells (Chernyakov, SantiagoV Tirado, and Bretscher 2013; Marston 2014) while at the same time damaged proteins, organelles and other aging factors are asymmetrically retained in the mother cell, eventually leading to senescence (Lai et al. 2002; McFalineVFigueroa et al. 2011; Aguilaniu et al. 2003; Sinclair and Guarente 1997; Dillin, Gottschling, and Nystrom 2014). Mutations affecting these two processes could lead to different demographic outcomes. Aberrant production or distribution of an asymmetrically distributed factor could lead to premature aging, whereas the failure to produce or distribute a critical subVcellular component could lead to newly born cells dying.

We found that fecundity and mortality were ageVstructured in two clinical isolates. In the strain YJM693, older cohorts showed lower fecundity and higher mortality, indicating that YJM693 cells senesce prematurely at elevated temperatures. Whatever causes senescence in budding yeast must be asymmetrically retained in mother cells (Kennedy, Austriaco, and Guarente 1994). A number of molecular mechanisms are associated with senescence in good conditions in yeast including autonomously replicating circularized rDNA, damaged vacuoles and mitochondria, and the aggregation of damaged proteins (Lai et al. 2002; McFalineVFigueroa et al. 2011; Aguilaniu et al. 2003; Sinclair and Guarente 1997; Dillin, Gottschling, and Nystrom 2014), although some of these remain controversial (Fehrmann, Paoletti, et al. 2013). We speculate that the strain YJM693 contains a temperature sensitive mutation in a gene required for normal replicative lifespan.

In contrast to the strain YJM693, older cells in the strain YJM996 were more fecund than younger cells. The low fecundity of young cohorts in these strains is partly, but not entirely, due to very high mortality in young cohorts. In contrast, the mortality of older cohorts is low, leading them to accumulate in the population. The force of natural selection on fecundity of a cohort scales with the relative survivorship of that cohort (Hamilton 1966). This implies that natural selection on the fecundity of older cells is much stronger in this strain, but only during heat stress.

The early life mortality of YJM996 is linked to a defect in mitochondrial inheritance. YJM996 cells contain mitochondria with an aberrant globular morphology during heat stress, and daughter cells in this strain frequently fail to inherit mitochondria. Although mtDNA is dispensable in *S. cerevisiae*, the mitochondria themselves are essential (Kispal et al. 2005; Chernyakov, SantiagoV Tirado, and Bretscher 2013; Frederick, Okamoto, and Shaw 2008). This strongly suggests that the failure to inherit mitochondria causes the high mortality in young cells in this strain, although we cannot rule out that other essential components may fail to be segregated as well.

Genetic variation affects the relative contribution of mortality and fecundity to stressVinduced changes population growth rate. For example, increased mortality was both necessary and sufficient to explain the slow growth of the clinical isolate YJM693. Cells that survived chronic heat stress in this strain had fecundity identical to unstressed cells, whereas nearly all cells with low fecundity were dead. In contrast, cells with low fecundity during heat stress in the clinical isolate YJM996 or the genomic reference strain S288C are sometimes alive. Our results indicate that changes in population growth rate should be interpreted cautiously, since they can reflect the effects of distinct demographic mechanisms.

The ability to grow at high temperatures in the presence of alcohol is a key innovation critical to the ecological success of *Saccharomyces cerevisiae* (Goddard 2008). In this study, we examined the standing genetic variation present within *S. cerevisiae* for the ability to grow at high temperatures. The ability of cells to grow at high temperatures has been hypothesized to be a trait that enables pathogenesis in yeast, but we show that clinical isolates can also be sensitive to heat. Thermal tolerance in yeast is a genetically complex trait (Steinmetz et al. 2002; Sinha et al. 2006), and consistent with this we found that heat can operate on distinct demographic functions to lower growth rate, reflecting effects on different cellular processes involved in asymmetric division in budding yeast. Our results connect physiology to demography, and shed light on how genetic variation affects the thermal limits of budding yeast.

## Methods

All of our experiments were conducted using clonal populations grown in YPD or in SC-ura media when a plasmid was used. Before staining, cells were propagated for at least 16 hours in log-phase growth at the temperature being surveyed. Cells were first stained with Alexa488-conjugated wheat germ agglutinin (WGA) for 15 min. Cells were washed once in YPD and diluted to 0.33X10^6^ cells/ml, then grown for six hours. Cells were then fixed in PBS + 8% formaldehyde, and stored until imaging. Before imaging, cells were stained as above with tetramethylrhodamine (TMR)-conjugated WGA. Widefield Z-stacks were acquired, deconvolved, and then processed using a maximum pixel intensity projection. The numbers of buds stained by each dye were counted by manually from these projections. At least 150 fields of view which contained at least one cell with the first stain were collected for each biological replicate. This number was chosen to keep the acquisition time required for a biological replicate of a two-color experiment below one hour. Growth curves were collected on a Tecan Sunrise over a 48hr period with measurements every 15 minutes. See Supplementary Information for detailed methods including for three-color TrackScar.

## Acknowledgements

We acknowledge Nicolas Buchler, Daniel Lew, Daniel Skelly, and Helen Murphy for helpful comments on the manuscript. We thank Debra Murray for suggesting the name “TrackScar”. This work was supported in part by NIH (P50GM081883) and NSF (MCB1330545) awards to P. M. M.

## Author contributions

C.S.M designed the experiments, carried out the experiments, analyzed data, and prepared the manuscript. P.M.M. designed the experiments, and prepared the manuscript

## Materials and correspondence

Material requests and correspondence should be directed to Paul M. Magwene.

## Supplementary Information Table of Contents

### Supplementary Introduction

### Supplementary Results

TrackScar provides a sensitive measure for differences in fecundity

TrackScar minimally affects cellular physiology

Population growth rate and mean fecundity are well-correlated

Six hours of recovery is sufficient to distinguish live from dead cells

Heat stress can alter the distribution of ages in a population

### Supplementary Methods

TrackScar Staining

Microscopy

Image Processing and Data Analysis

Data Collection and Microscopy for Time Series

Growth Curves Strain Construction

### Supplementary References

### Supplementary Figures

*Figure S1.* TrackScar is sensitive and minimally affects cellular physiology

*Figure S2.* Correlation between TrackScar and growth curves

*Figure S3.* Growth rate of all strains as a function of temperature

*Figure S4.* Histograms of all strains analyzed using TrackScar

*Figure S5.* Age-structured fecundity of S288C

*Figure S6.* Distribution of ages during heat stress

*Figure S7* Penetrance of globular morphology phenotype

*Figure S8* Fecundity of cells with globular mitochondria

### Supplementary Tables

*Table S1.* Strains used in this study

## Supplementary Results

### TrackScar minimally affects cellular physiology

A potential concern with any vital-staining method is the possibility that it could change the cell physiological processes of interest. To explore this possibility, we compared division time estimates based on TrackScar to results from previous studies using timelapse microscopy. Using TrackScar we estimated the average division time to be 73.9 minutes for haploid cells of the genomic reference strain S288c grown in rich-media conditions. This is very similar to an estimate of 72.6 minutes derived previously for this same strain using timelapse microscopy (Lord and Wheals 1981). To further explore the physiological impact of WGA staining on cellular division rates, we carried out a TrackScar experiment in which the time interval between the first and second stain was varied between one and six hours. We reasoned that if WGA staining is a significant cellular stress that the reproductive rates immediately following the first stain should be systematically lower than in later time points, because extended incubation would allow cells to recover from any physiological perturbations caused by staining. We carried out this experiment for nine genetically diverse *S. cerevisiae* strains. We found no evidence that reproductive rates at earlier time points were any lower than later time points (Fig. S1a). Indeed, our data show that cells at time points immediately following the first stain produce slightly more daughters than those at later time points (linear model; p = 0.03). Taken together, these two sets of comparisons suggest that TrackScar does not significantly perturb the division rate of *S. cerevisiae cells.*

### Trackscar provides a sensitive measure of differences in fecundity

To determine the sensitivity of TrackScar, we tested whether this method could detect the increased cell cycle time of daughter cells. *S. cerevisiae* reproduces by asymmetrical division, and smaller daughter cells must grow longer until they reach a critical size threshold before dividing for the first time (Hartwell and Unger 1977). Thus, daughter cells should have lower fecundity than mother cells that had already divided at least once. Consistent with this expectation, daughter cells of haploid strain S288C produced an average of 4.9 daughters in a six-hour period, whereas mother cells produced an average of 5.4 daughters (Fig. S1b). This difference is significant (Paired t-test; n=3; p = 0.030). These results indicate that TrackScar can accurately detect small differences in fecundity between individual cells.

### Population Growth Rate and Mean Fecundity Are Well Correlated

The different growth rates of sensitive and robust strains provided another opportunity to test the accuracy of TrackScar. We used two-color TrackScar to estimate fecundity at 30C and 35.5C for 8 sensitive and 13 robust strains. We compared these estimates of average fecundity to the estimates of growth rate obtained from growth curves. The average fecundity of cells measured using TrackScar and the maximum population growth rate measured by optical density at 35.5C are well-correlated (r^2^ = 0.58; Fig. S2).

### Six Hours of Recovery Is Sufficient To Distinguish Live from Dead Cells

Three-Color TrackScar relies on a sufficiently long recovery phase to distinguish between live and dead cells. We tested whether six hours is sufficient to distinguish live from dead cells by determining if YJM693 cells that did not divide after six hours of recovery would subsequently form a colony on an agar plate.

We grew YJM693 to early-log phase at 35.5C as and stained the cells with Ax488-WGA as described in the methods for the TrackScar protocol. After washing off the stain, we spread the cells on a plate of YPD and randomly selected cells from the patch of cells on the plate. Cells were immediately singled and subsequently moved into a grid on the YPD plate. If a cell divided in between the time that the cell was singled and when it was moved to the grid, both the mother cell and its progeny were placed in the same space on the grid. The plate was incubated for 6hr at 30C and colonies were inspected for new daughter cells using a microscope. After 48hr, the number of colonies formed by founder cells was recorded. Out of 24 cells that did not divide after six hours of recovery at 30, only one of these these cells subsequently divided in the next 48 hours. This indicates that six hours of recovery in good conditions is adequate to distinguish live cells from dead cells.

### Heat stress can alter the distribution of ages in a population

In an exponentially growing yeast population, the frequency of cells of age n in the population would be approximately 1/2^*n*^ if age had no influence on fecundity or mortality (the asymmetric division of yeast leads to a more complicated actual distribution of ages that approximates 1/2^*n*^ when the doubling time of the population is short (Lord and Wheals 1980)). Therefore, without specifically enriching for old cells, it should be quite rare to observe cells that have budded more than about ten times as they should make up less than 1 in 1,000 cells in the population. However, as described above, heat stress can induce changes in age related patterns of fecundity and mortality. One would predict therefore that some stress related changes, particularly those that deplete young cells, could be sufficient to alter population demography broadly.

We found that in the absence of heat stress, the distribution of ages in the populations of all three strains approximately matched the 1/2^*n*^ expectation, consistent with no influence of age. Furthermore, neither YJM693 nor S288C showed significantly different age distributions at 30°C and 35.5°C (Kolmogorov-Smirnov test, p > 0.3). However, YJM996 had a significantly different distribution of ages during growth at 35.5°C (Kolmogorov-Smirnov test, p = 1.74X10^−7^). This difference is due to a strong enrichment of cells older than five buds and a depletion of cells between 2 and 4 buds (Fig. S6). Consequently, we frequently found cells with more than 15 buds in YJM996 populations during growth at 35.5°C. This is consistent with the low fecundity and increased mortality we observed in young cells and indicates that heat stress can substantially alter the age distribution of a population of cells by affecting their demographic rates.

## Supplementary Methods

### TrackScar Staining

All of our experiments were conducted using recently clonal cell lines in early-to mid-log-phase growth, grown in YPD media (Sherman 2002), except for experiments to visualize mitochondrial morphology, which where conducted in synthetic media lacking uracil with 2% dextrose (SC-ura) media. To begin each experiment, frozen stocks were streaked onto YPD or SC-ura plates to obtain single colonies and incubated at 30°C. After 24 to 48 hrs, single colonies were inoculated into 2ml of YPD or SC-ura overnight at 30°C.

To obtain log-phase cells, overnight cultures were diluted 1:100 and then in a 1:4 serial dilution in the columns of a 96 well plate. The final volume in each well of the plate was 150p! The diluted cells were then placed in an incubator and shaken at 500rpm at the desired temperature for at least 16 hr. Dilutions containing putative early log-phase cultures were located by examining the optical density of cells by eye and were then counted in a hemocytometer. A dilution whose concentration was between 0.7×10^6^ and 7×10^6^ cells/ml was chosen.

Log-phase cells were gently spun out of solution by centrifugation at 500xg for 1.5 minutes. All centrifuge steps on live cells were done at this speed. Cells were stained in 60μl of fresh media with 33μg/ml wheat germ agglutinin (WGA) conjugated to Alexa488 (Invitrogen, W11261) at 30C for 15 min on a rotor. Cells were washed once in 200μl of fresh media then diluted to 0.33×10^6^ cells/ml in one column of a 96 well plate. Cells were then placed back in the incubator for 6 hr.

To fix the cells for subsequent imaging, cells were spun out of solution at 5,000xg for 2 minutes. All subsequent centrifugation steps were done at this speed. Cells were resuspended in 500μ.l of sterile PBS. Cells were fixed by adding 100μ.l of 37% formaldehyde to the PBS and incubating at room temperature for at least 15 minutes. Cells were washed once in sterile PBS then resuspended in sterile PBS and stored at 4°C in the dark until imaging. Cells were generally imaged less than a week after collection, but the staining was stable for at least three months.

Immediately prior to imaging, cells were spun out of PBS and resuspended in 60#x03BC;.l of fresh media with WGA conjugated with tetramethylrhodamine (Invitrogen, W849) for 15 minutes at 30°C on a rotor. Cells were washed 1× in 500μl of sterile PBS, then sonicated in six short bursts using a Branson Sonifier 150 at setting 4 to break up any clumps.

For experiments examining mitochondrial morphology, TrackScar staining was performed as above, except the first WGA stain was WGA-TMR and the second WGA stain six hours later was WGA-640CF (Biotium).

For the “Three-Color TrackScar” experiments used to determine if cells with low fecundity during heat stress would divide subsequently after incubation at 30C, cells were: 1) stained as above with WGA-Ax488 at the beginning of the experiment; 2) stained as above with WGA-640CF after 6hr of incubation at either 35.5°C or 40°C; 3) stored overnight at 4°C in YPD (for experimental convenience); 4) incubated in YPD for 6hr at 30C on a rotor drum; 5) fixed with formaldehyde as above and stored, and; 6) stained as above with WGA-TMR.

### Microscopy

To count the stained bud scars for all experiments except for the time series in Fig. S1 (see below), cells were imaged on a DeltaVision wide-field fluorescent microscope using a 100x lens. Since cells that do not have the first stain (WGA-Ax488) do not have any information about the number of daughters produced by the cell, we randomly imaged cells that were stained with the first stain. To find cells to image, we moved in transects along the slide and imaged each cell that had staining in the Ax488 channel. The exception to this was when we sought out old cells specifically in order to measure the fecundity or mortality of older cohorts. In this case, we counted the number of buds that were visible in the Ax488 channel and ignored young cells.

For each field of view containing a cell, ten-to twelve-micron deep z-stacks with 0.2 to 0.4 micron slices were collected at every point with a 0.1 second exposure with filter sets appropriate for each fluorophore being used in that experiment. The depth of the z-stacks was chosen to ensure that both sides of every cell were imaged. To visualize cell outlines, z-stacks of Nomarski bright-field images were collected with a 0.025 second exposure. Either 768×768 or 1024x1024 images were collected. Z-stacks were deconvolved using the default settings of the DeltaVision software. The deconvolved z-stacks were cropped to remove deconvolution artifacts. Each channel was projected onto a single slice using a maximum pixel intensity projection. At least 150 images (containing at least one cell) for each sample were collected.

To generate images for display items in the figures, the brightness and contrast of each cell and each channel was adjusted independently so that the first stain would be visible. The gamma of the brightfield channel was sometimes adjusted in order to remove saturated pixels that made display of the brightfield image difficult.

### Image processing and data analysis

The projected images of each z-stack were assembled into TIFF stacks using a custom Python script and ImageMagick. Individual cells with the first TrackScar stain (either Ax488 or TMR) were cropped from the wider images using custom ImageJ macros. To count the bud scars, a false-color image of the individual cell was displayed. To aid in the counting, we wrote ImageJ macros that allowed the rapid toggling of different channels. Each bud scar in each channel was counted using the following rules: 1) the birth scar is always counted as the first scar; 2) if a bud scar has any staining of the previous stain that is visible at a normal brightness and contrast level, then that bud scar is counted as being stained by that stain; 3) bud scars that correspond to daughter cells still attached to the mother cell are not counted unless the daughter cell has divided (has a bud scar besides its birth scar).

Mitochondrial morphology for Fig. S7 was scored by examining at least 100 random cells for each biological replicate in an Axio Imager and using a 100x lens.

The field aperture was adjusted so that only one cell was visible at a time, and each cell on a transect was scored. The presence of clumps of mitochondria and mitochondrial threads was scored by examining cells using the appropriate filterset for GFP. Mitochondrial morphology for Fig. S8 was scored by examining maximum projections of Z-stacks created using a DeltaVision microscope as above for at least 150 cells for each biological replicate. The presence of clumps of mitochondria and mitochondrial threads was scored before examining the TrackScar staining for each cell.

All statistical tests were performed using R (R Core Team 2015). To estimate the division rate of haploid S288C cells, we fit a linear model to a plot of buds produced as a function of time. To test for the effect of age on the average fecundity of a cohort, we computed the average fecundity of the cohort of each strain for three biological replicates and fit a linear model to those averages.

### Data collection and microscopy for time series

To collect the time series in Fig. S1, we used the diploid strains YJM1529, YJM1549, YJM1573, YJM195, YJM554, YJM555, YJM693, and S288C as well as haploid S288C cells. TrackScar staining was done as above except the time between the two stains was varied between one and six hours. To count the number of daughters produced by cells in the time series in Fig. S1, the number of buds on at least fifteen random cells stained by each stain were counted were counted using an Axio Imager and a 100x lens in “real time” (without acquiring images). To see buds on both sides of the cell, the plane of focus was moved up and down.

### Growth curves

To measure the maximum population growth rate of strains in the 100 genomes collection, 93 strains (see Table 1) were grown overnight in a 96 well plate with 100μl of YPD per well at the desired temperature on a plate shaker at 280rpm. Cultures were mixed and diluted 1:5000 into a 96 well plate with 100μl of YPD per well. Growth curves were measured in a Tecan Sunrise plate reader. Measurements were collected every 15min for 48hr with 30s of shaking at the highest setting before each measurement. Four biological replicates were collected for the temperatures 30°C and 35.5°C and at least two replicates were collected for all other temperatures.

The maximum growth rate of a culture was calculated using non-parametric smoothing using the “cellGrowth” package from Bioconductor. To identify strains sensitive to heat stress, we looked for strains whose maximum population growth rate at 35.5°C was 93% or lower than their growth rate at 30°C. This cutoff was chosen because S288C populations grew 94% as quickly at 35.5°C at 30°C.

### Strain construction

To generate the uracil auxotrophs CMY145 and CMY146 in the YJM693 and YJM996 genetic background, respectively, the URA3 gene was disrupted by a KanMX3 cassette. The plasmid M3927 (Voth, Wei Jiang, and Stillman 2003) (aka PMB161) was digested using BamHI, then transformed into YJM693 and YJM996 using LiAc/PEG transformation (Gietz and Schiestl 2007). G418 resistant colonies were checked for the insertion of the cassette using colony PCR and strains with the insertion of the cassette were sporulated. Tetrads were dissected and G418 resistant, uracil auxotrophs were identified by replica plating. To visualize mitochondrial morphology, the plasmid p416-GPD-mito-roGFP1 (aka PMB273) (McFaline-Figueroa et al. 2011)(a kind gift from Liza Pon) was transformed into CMY145, CMY146, and PMY044 (a uracil auxotroph in the S288C background) to yield CMY177, CMY178, and CMY182, respectively.

## Supplementary Figures

**Figure S1.**
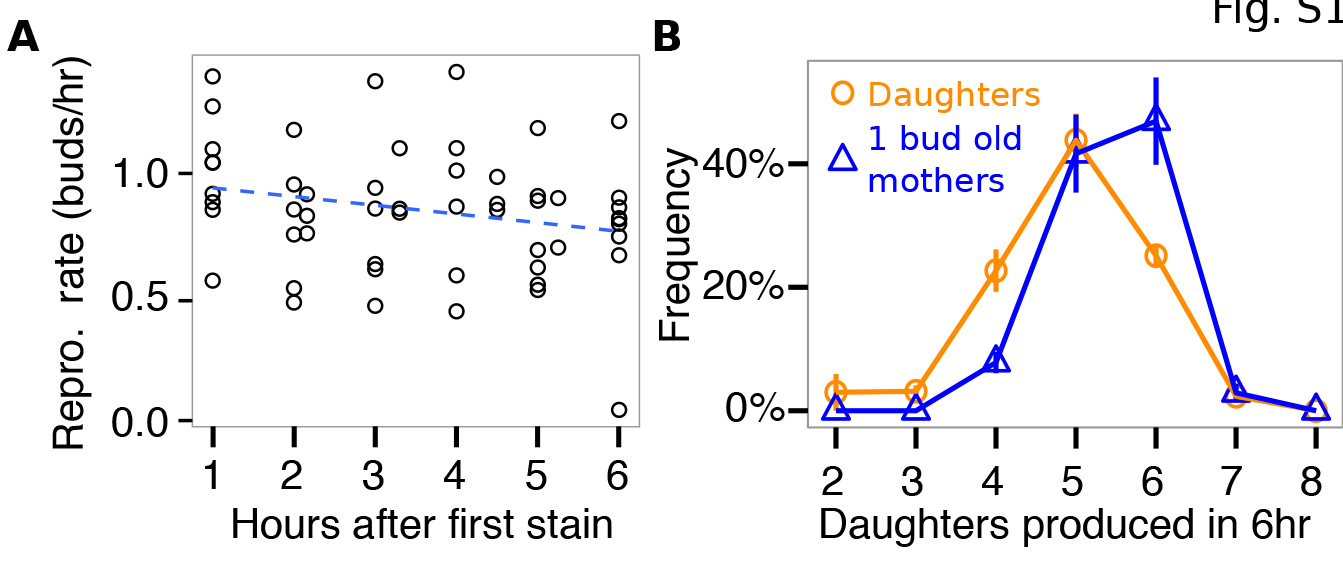
TrackScar is sensitive and minimally affects cellular physiology. **A)** The average number of daughters produced in a one hour interval by cells in nine genetically diverse yeast strains is shown as a function of time since the first WGA stain. A linear regression line is plotted as a blue dashed line. **B)** Histograms of the number of daughters produced by daughter cells (orange circles) and mother cells that had divided once (blue triangles) in a 6hr interval for are shown. Error bars show the standard error of the mean for three biological replicates.Mean division rate (daughters per hour)

**Figure S2.**
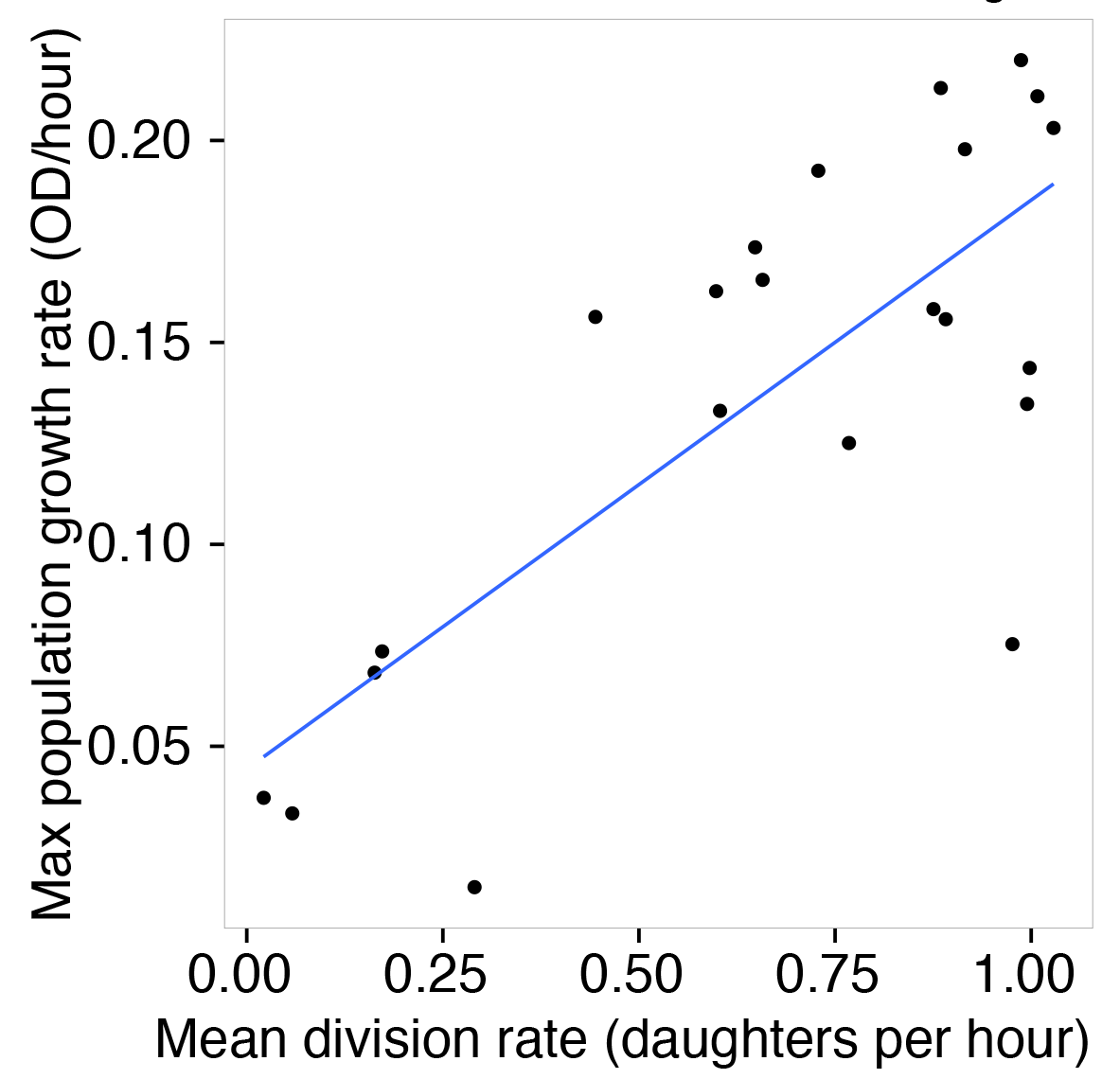
Estimates of the growth rate of a population made using TrackScar and growth curves are well-correlated. A scatter plot of the max growth of a population inferred by measuring its rate of increase in a spectrophotometer and the average fecundity of a population measured using TrackScar is shown for a diverse panel of yeast strains grown at 35.5C. A linear regression line is shown in blue.

**Figure S3.**
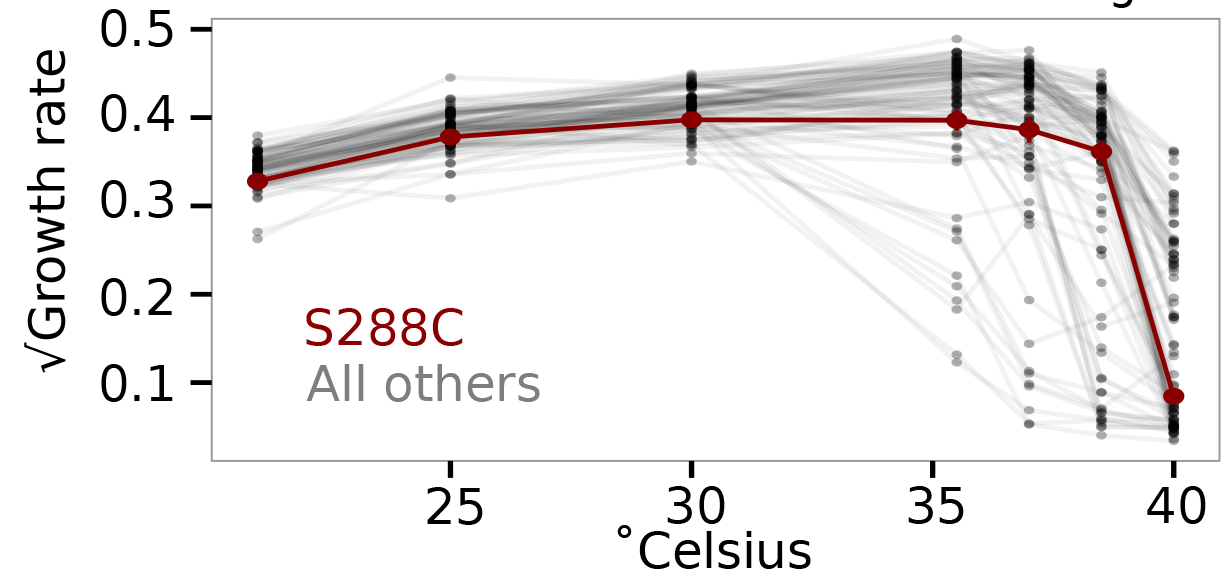
There is extensive genetic diversity in thermotolerance among environmental yeast isolates. The square-root of the maximum growth of each of 93 yeast strains is shown as a function of temperature. The strain S288C is shown in red.

**Figure S4.**
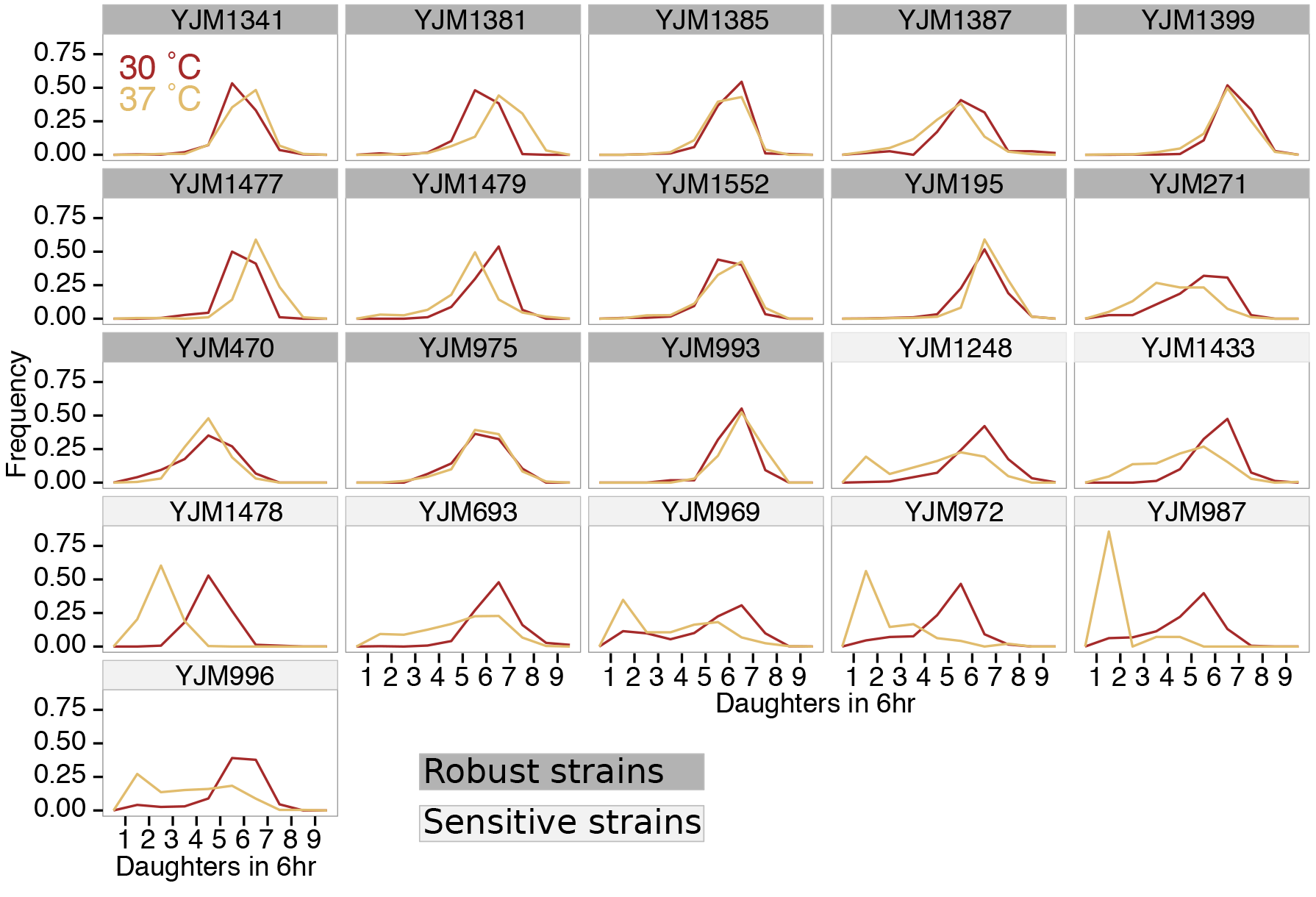
TSensitive strains frequently show sub-populations of cells with very low fecundity. Plots showing histograms of the number of daughters produced in 6hr at 30C (brown) and 35.5C (gold) are shown for all strains that were examined using TrackScar. Strains that are robust to heat stress have their facet labels colored in dark grey.

**Figure S5.**
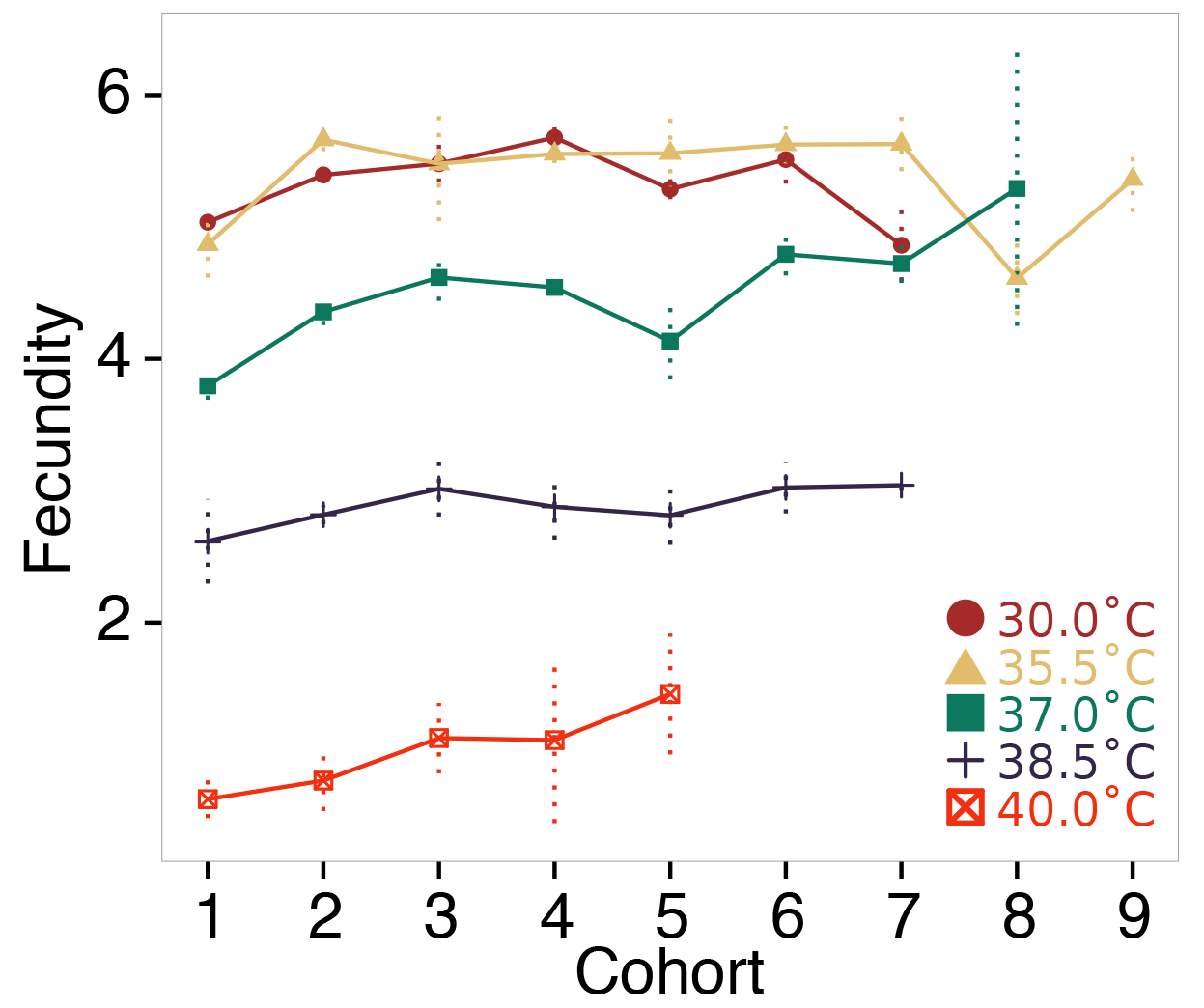
The fecundity of S288C cells is not age structured at any temperature. The average fecundity of cohorts of S288C cells during 6hr during growth at 30C (brown circles), 35.5C (gold triangles), 37C (green squares), 38.5C (black crosses), and 40C (red crossed boxes). Error bars are the standard error of the mean for at least two biological replicates. Only cohorts with at least three cells in each of the biological replicates are shown

**Figure S6.**
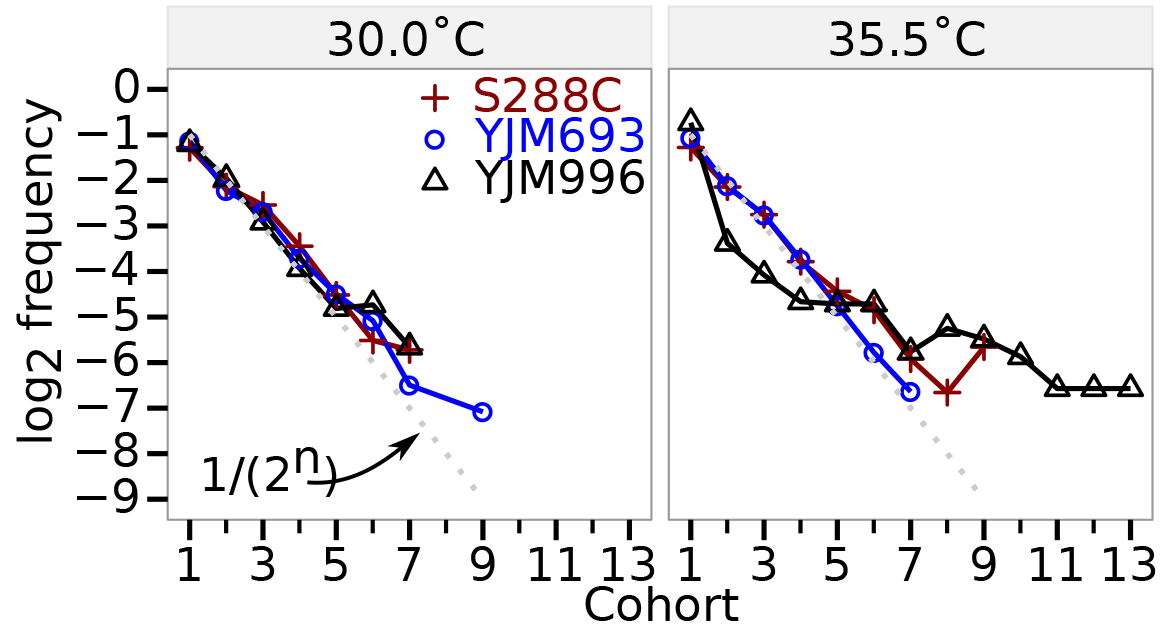
Heat stress can alter the distribution of ages in a population. The frequency of cells in a cohort in a sample of randomly chosen cells at 30C and 35.5C is shown for the strains S288C (red crosses), YJM693 (blue circles), and YJM996 (black triangles) is shown. The grey dotted line shows 1/(2^*n*^). Each point represents the pooled observations across at least three biological replicates. Only cohorts with at least five total cells are shown.

**Figure S7.**
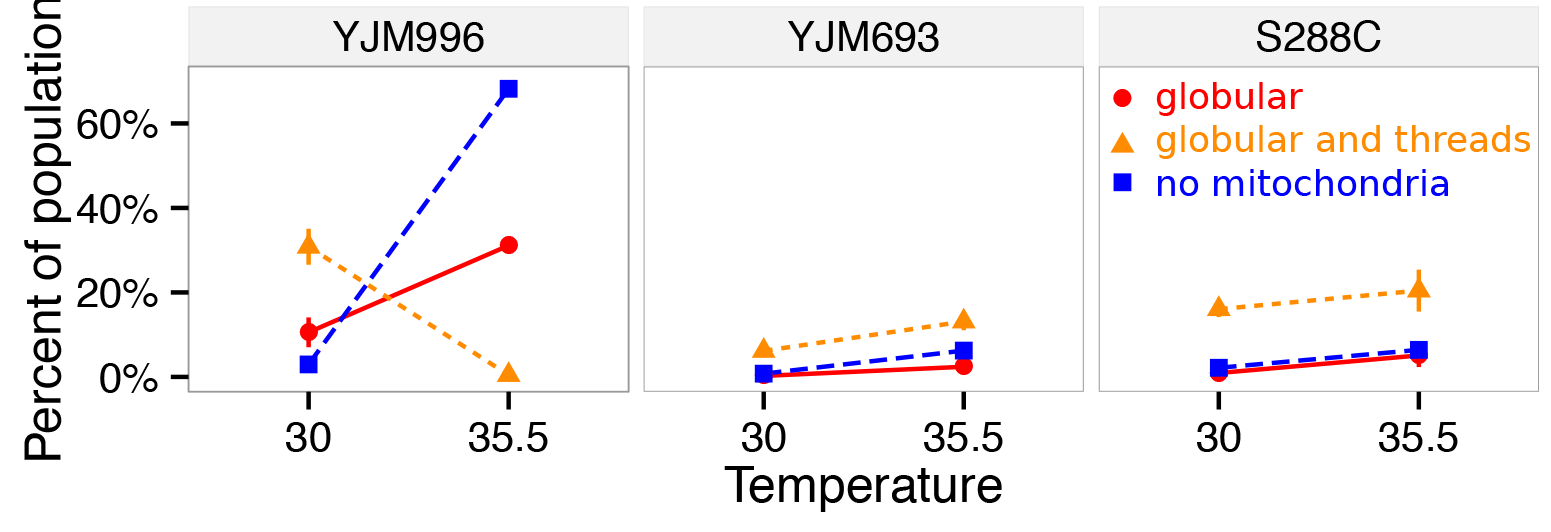
Heat stress changes the frequency of cells with abnormal mitochondrial morphology in the strain YJM996. The percentage of the population with abnormal mitochondria at 30°C and 35.5°C is shown for the strains CMY177 (YJM693 genetic background), CMY178 (YJM996 genetic background), and CMY182 (S288C genetic background). Mitochondria were grouped into the classes: globular (red circles), globular with some threads (orange triangles), and no mitochondria (blue squares). Error bars show the standard error of the mean for three biological replicates of at least 100 cells.

### Supplementary Tables

**Supp. Table 1**Strains used in this study. See attached file “TableSl - strains.xlsx”

